# Decoding the regulatory logic of the *Drosophila* male stem cell system

**DOI:** 10.1101/227876

**Authors:** Fani Papagiannouli, Srividya Tamirisa, Eugen Rempel, Olga Ermakova, Nils Trost, Jun Zhou, Juliane Mundorf, Samantha Brunel, Naima Ruhland, Michael Boutros, Jan U. Lohmann, Ingrid Lohmann

## Abstract

In the past decade, the importance of the niche to provide regulatory inputs to balance stem cell self-renewal and differentiation has become clear. However, the regulatory interplay between stem cells and their niche at the whole genome level is still poorly understood. To elucidate the mechanisms controlling stem cells and their progenies as they progress through their developmental program at the transcriptional level, we recorded the regulatory program of two independent cell lineages in the *Drosophila* testis stem cell model. To this end, we identified genes active in the soma or germline as well as genome-wide binding profiles of two essential transcription factors, Zfh-1 and Abd-A, expressed in somatic support cells and crucial for fate acquisition of both cell lineages. Our data identified key roles for TOR signalling, signal processing V-ATPase proton pumps and the nuclear transport engaged nucleoporins and we demonstrate their importance in controlling germline maintenance, proliferation and differentiation from the support side. To make our dataset publicly available and support quick and intuitive data mining, we generated an interactive online analysis tool. Applying our tool for comparative analysis, we uncovered conserved core gene sets of adult stem cells across species boundaries. We have tested the functional relevance of these genes in the *Drosophila* testis and intestine and find a striking overrepresentation of stem cell defects when the corresponding genes were depleted. In summary, our dataset and interactive platform represents a versatile tool for identifying novel gene networks active in diverse stem cell types and provides a valuable resource for elucidating the multifaceted regulatory inputs required to guide proper stem cell behaviour.

## INTRODUCTION

Animal tissues and organs are generated and maintained by adult stem cells, which are maintained in an undifferentiated and proliferative state, while at the same time generating lineage-restricted daughter cells that undergo terminal differentiation. Due to this unique ability, they are able to replace dying, lost or damaged cells and consequently are critical for tissue homeostasis. Mis-regulation of this process can result in tissue degeneration or tumorigenesis. Thus, the balance between self-renewal and differentiation is tightly controlled and stem cell intrinsic mechanisms are known to play an important role in this process (Amoyel et al., 2013; Biteau et al., 2011; Pearson and Sánchez Alvarado, 2008). In addition, signals emanating from the local tissue environment, the stem cell niche, have been identified more recently to be equally important to control the activity of stem cells and their progeny (Hsu and Fuchs, 2012; Kitadate and Kobayashi, 2010; Martinez-Agosto et al., 2007; Morrison and Spradling, 2008). However, it is still poorly understood how niche cells execute their regulatory function on a global mechanistic level.

The *Drosophila* testis represents an excellent model for studying stem cell - niche interaction at the cellular and molecular level (de Cuevas and Matunis, 2011; Dominado et al., 2016). At the tip of the testis two types of stem cells are found, the germline stem cells (GSCs) and the somatic cyst stem cells (CySCs), which maintain spermatogenesis (Fuller and Spradling, 2007; Li et al., 2014). Each GSC is enclosed by two CySCs and both stem cell types are anchored to a group of eight to 16 nondividing somatic cells, called the hub (Fairchild et al., 2015; Hardy et al., 1979), through adhesion molecules (Matunis et al., 1997; Wang et al., 2006). GSCs and CySCs divide asymmetrically to produce daughter cells, gonialblasts (Gbs) and somatic cyst cells (SCCs), respectively that form a developmental unit called cyst. Whereas gonialblasts continue to divide and undergo four rounds of mitotic divisions to produce 16 spermatogonia that further develop into spermatocytes, SCCs grow without division and co-differentiate with the germline (Figure 1A) (Cherry and Matunis, 2010; Fuller and Spradling, 2007).

**Figure 1.**
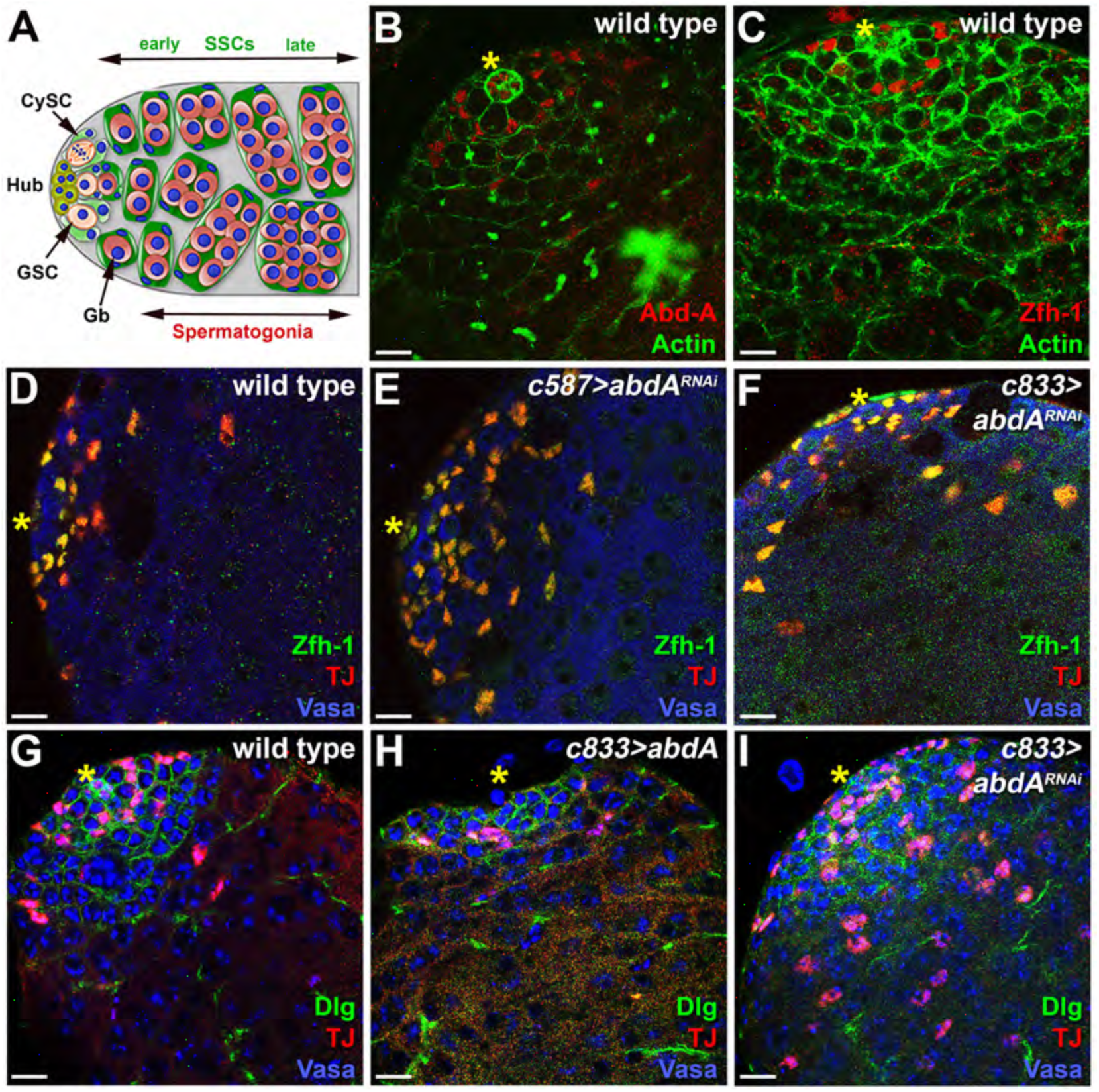
The Hox transcription factor Abd-A controls the switch from early to late SCCs. **(A)** Schematic diagram of early spermatogenesis in *Drosophila* 3^rd^ instar (L3) larval testis. GSC, germline stem cell; CySC, somatic cyst stem cell; Gb, gonialblast; SCCs, somatic cyst cells. **(B, C)** L3 wild-type testes stained for Abd-A (red) (B) or Zfh-1 (red) (C) and Actin (green) to mark cyst cells and germline fusomes. **(D-F)** L3 wild-type (D), *c587*>*abdA^RNAi^* (E), *c833*>*abdA^RNAi^* (F) testes stained with TJ (red), Zfh-1 (green) and Vasa (blue). Knock-down of *abd-A* in cyst cells results in expansion of Zfh-1 expression, resulting in its co-localization with TJ in CySCs and early SCCs. **(G-I)** L3 wild-type (G), *c833*>*abdA* (H), *c833*>*abdA^RNAi^* (I) testes stained for TJ (red), Dlg (green) to mark the hub and cyst cells and DAPI for the DNA (blue). Overexpression of *abd-A* in cyst cells results in reduced numbers of TJ positive cyst cells (H) while the number is increased when *abd-A* levels are reduced by RNAi (I). Yellow asterisks mark the location of the hub. Testes are oriented anterior left. Scale bars, 10 μm. See also Figure S1.

The somatic cell population fulfils several important support functions in the *Drosophila* testis. On the one hand, somatically derived hub cells express the growth factor Unpaired (Upd), the ligand that activates the JAK-STAT pathway in the adjacent stem cells, thereby instructing CySCs maintenance (Kiger et al., 2001; Matunis et al., 1997; Tulina and Matunis, 2001) and regulating GSC anchoring to the hub (Leatherman and DiNardo, 2008; Stine et al., 2014). On the other hand, CySCs and their progenies seem to be also critically required for the development of the germline. Intriguingly, CySCs not only induce GSC fate by signalling to the neighbouring germline cells via the Transforming Growth Factor-β (TGFβ) pathway (Kawase et al., 2004; Shivdasani and Ingham, 2003; Slaidina and Lehmann, 2014), but CySCs also instruct the self-renewal of GSCs cells in a cell non-autonomous manner. One factor, the transcriptional regulator Zinc-finger homeodomain protein 1 (Zfh-1), which is activated by hub-mediated JAK-STAT signalling in CySCs, is key for the soma-germline crosstalk as well as for CySC development (Leatherman and DiNardo, 2008; Llorens-Bobadilla et al., 2015; Signer et al., 2014). Importantly, Zfh-1 alone is sufficient for the induction of GSC self-renewal, even outside the niche (Leatherman and DiNardo, 2008). Zfh-1 controls a so far uncharacterized gene network in the soma essential for soma and GSC maintenance.

Close communication between somatic and germline derived cells is not only required for germline stem cell maintenance but also for later stages of spermatogenesis. For example, survival of the germline specifically at the spermatocyte stage depends on Eyes absent (Eya) and Sine oculis (So) (Fabrizio et al., 2003; Kiger et al., 2001), again two transcription factors (TFs) active exclusively in late SCCs. Testis homeostasis is not only dependent on the soma-germline signalling, but also requires the direct physical contact of both cell populations. Consequently, spermatogonia as well as their enclosing SCCs die in mutants of the tumor suppressor *discs large* (*dlg*), a septate junction protein (Leatherman and DiNardo, 2008; Papagiannouli and Mechler, 2009). All these studies highlight the importance of the soma for the integrity and activity of the niche and stem cells, as well as for the coordinated maintenance of the testis as an entity. Despite its crucial function not only in the *Drosophila* testis but also in other stem cell systems, our knowledge of how support cells control the behaviour of stem cells and their progenies is still very limited.

Here, we applied genome-wide *in vivo* mapping of DNA-protein interactions to comprehensively determine genes active in the *Drosophila* larval testis soma and germline as well as genes targeted by two key TFs with critical roles in the somatic lineage. One of them is the known stem cell maintenance factor Zinc finger homeodomain 1 (Zfh-1) ( Kiger et al., 2001; Leatherman and DiNardo, 2008; Sinden et al., 2012), while the other is the Hox TF Abdominal-A (Abd-A), for which we uncovered a novel stem cell regulatory function in the *Drosophila* testis. Meta-analysis of these datasets not only recovered the core group of genes known to control stem cell development in the *Drosophila* testis, but identified a large number of novel regulators with critical functions. To make our dataset easily accessible and amenable to comparative analysis using other datasets, we have developed an interactive and versatile data mining and analysis tool, which is openly available online. Extensive data cross-comparison using this tool allowed us to identify the major processes active in the soma. Using interaction network approaches and transgenic RNAi, we found that selected gene networks are critical for testis development. Furthermore, by comparing our data to published datasets from diverse stem cell systems and organisms, we uncovered a core gene expression set of adult stem cells conserved across species boundaries. Testing TFs from this category by RNAi, we find an unusually high frequency of drastic stem cell phenotypes in two *Drosophila* stem cell systems, suggesting that the core expression set defines regulators with substantial functional relevance. Moreover, our analysis also revealed system specific regulators, which may represent the factors that allow individual cell types to respond to and interact with their typical yet diverse microenvironments. Taken together, we not only elucidate some of the mechanisms used by somatic support cells to control the activity of neighbouring germ cells, but also provide a rich resource to identify further important (and conserved) regulators of the balance between proliferation and differentiation in a genetically tractable stem cell system.

## RESULTS

### Defining the Hox transcription factor Abd-A as a novel regulator in the *Drosophila* testis soma

To study how the progressively maturing support cells control their own and the behaviour of the closely adjoining and co-differentiating germline, we aimed at comprehensively elucidating gene activities in the somatic lineage (CySCs, early SCCs, late SCCs). Unfortunately, cell-sorting approaches are so far not feasible using the *Drosophila* testis due to limitation in cell number and interference from tissue adhesiveness. Similarly, cell type specific targeted gene expression profiling cannot be applied due to the unavailability of stem cell- and cyst cell-specific regional cisregulatory regions. To circumvent these limitations, we used an alternative strategy: We first determined the transcriptome of the somatic and germline lineages by RNA polymerase II Targeted DamID (TaDa) (Kiger et al., 2001; Southall et al., 2013; Tulina and Matunis, 2001), respectively to identify and discriminate soma- and germline-specific expression profiles. In a next step, we elucidated the genes bound by two key regulators exclusively active in cells of somatic sub-populations and critically controlling their development using regular DNA adenine methyltransferase identification (DamID) (Kitadate and Kobayashi, 2010; van Steensel et al., 2001). By combinatorial analysis of these datasets we expected to identify a substantial fraction of genes specifically active in *Drosophila* testis support cells and thus contributing to their cell-autonomous and non-autonomous functions.

The first regulators we employed for our profiling was the well-described TF Zfh-1, which is highly expressed in CySCs and at lower levels in their immediate daughter SCCs (Figures 1C, 1D, S1B). Zfh-1 blocks differentiation of CySCs and indirectly controls GSC self-renewal from the soma (Amoyel et al., 2013; Leatherman and DiNardo, 2008; Michel et al., 2012). However, Zfh-1 has no major role in early SCCs, a cell population that supports the first stages of germline differentiation. In order to elucidate the genes actively regulating these events within early SCCs, we aimed at identifying another TF expressed in these cells and controlling their behaviour. To this end, we screened our whole-soma transcriptome dataset for TFs with known master-regulatory functions. One prominent candidate was the Hox TF Abd-A and protein expression analysis revealed that Abd-A accumulates in hub cells as well as in CySCs and early SCCs of 3^rd^ instar larval (L3) testes (Figures 1B, S1A), in a pattern similar to the known CySC and early SCC marker Traffic Jam (TJ) (Figures 1D, S1A, S1G) (Amoyel et al., 2016; Li et al., 2003). While Abd-A and Zfh-1 expression overlaps in CySCs and their immediate daughter cells (Figures 1A, 1B), they have clearly distinct functions in the testis. Cell-type specific knock-down of *abd-A* using two drivers active throughout the somatic lineage (except for the hub), *c587*-GAL4 (Figure 2A) (Li et al., 2007; Manseau et al., 1997) and *c833*-GAL4 (Figure S2A) (Amoyel et al., 2014; Papagiannouli and Mechler, 2009), resulted in an expansion of the number of CySC daughter cells, the early SCCs, co-stained with TJ and Zfh-1 in L3 testes (Figures 1D, 1E, 1F). Consequently, early SCCs were found in close association with late SCCs surrounding early spermatocytes (Figures S1J, S1K), which was never the case in wildtype L3 testes (Figure S1G). In contrast, knock-down of *zfh-1* using the *c833*-GAL4 driver interfered with the maintenance - differentiation balance of CySCs: the number of early SCCs expressing TJ was reduced (Figures S1G, 1I), while late SCCs labelled with Eya were observed in close proximity to the hub (Figures S1D, 1F), which is similar to the effects of soma-specific Abd-A over-expression (Figures 1G, 1H, S1E, S1H). These results led us to hypothesize that Abd-A regulates the identity of early and late cyst cells and that Abd-A levels are critical for the switch from CySC to early SCC and then to late SCC fate, while Zfh-1 functions primarily in CySCs (Buchon et al., 2009; Klimmeck et al., 2012; Leatherman and DiNardo, 2008). Interestingly, our analysis revealed a regulatory and functional interaction between the two TFs: Abd-A interacted with genomic regions in the vicinity of the *zfh-1* gene (Figure 2E), and Zfh-1 expression was expanded after Abd-A knock-down leading to a co-localization with the CySC and early SCC marker TJ (Figures 1D, 1E, 1F). Importantly, the expansion of Zfh-1 expression in response to reduction of Abd-A function did not induce the massive over-proliferation of somatic and germline populations reported for Zfh-1 overexpression (Leatherman and DiNardo, 2008).

**Figure 2.**
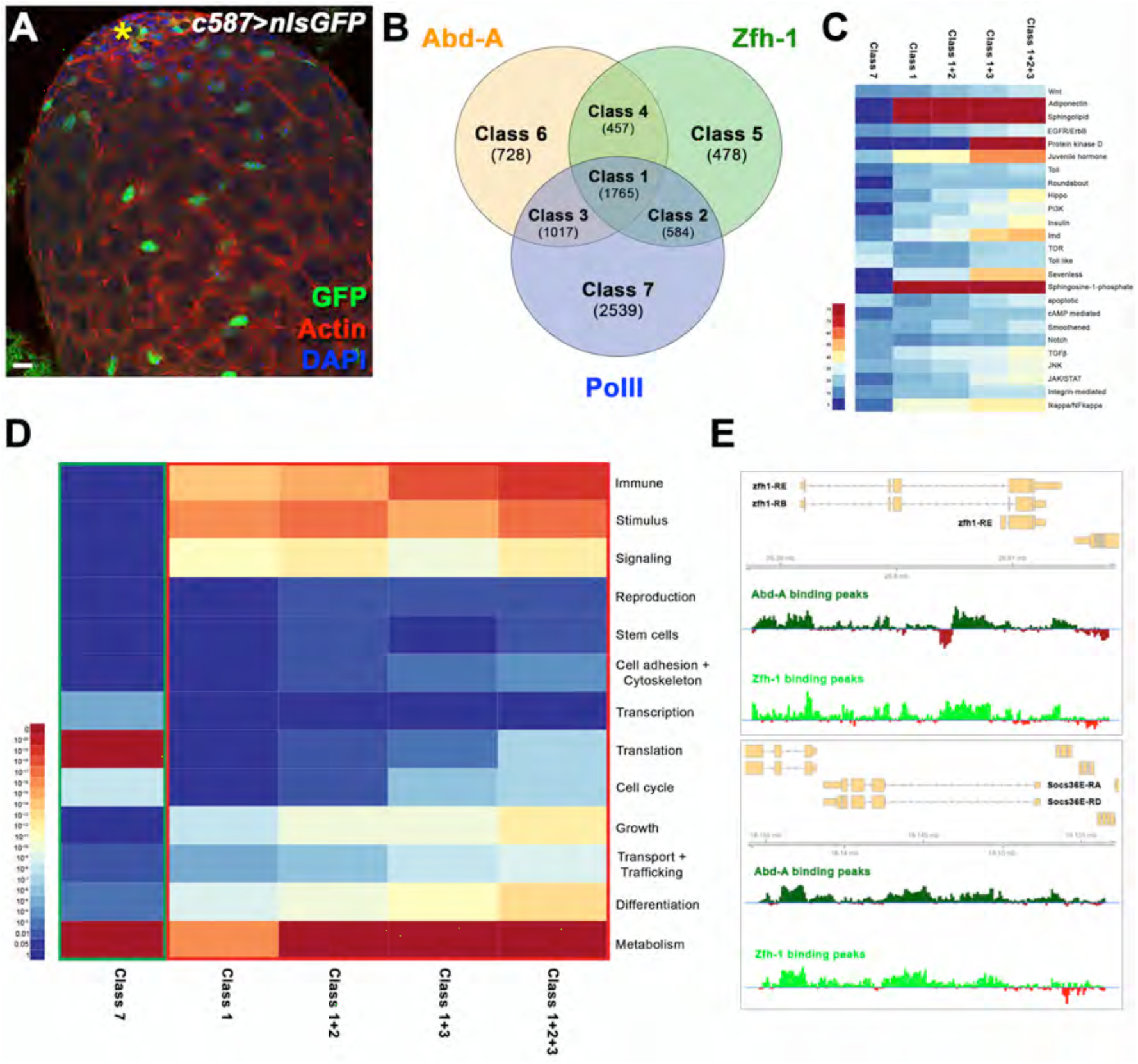
Profiling and comprehensive analysis of the *Drosophila* testis soma. **(A)** *c587*>*nlsGFP* L3 testis stained with GFP, Actin to mark cyst cells and DAPI for the DNA, highlighting the expression of the *c587*-GAL4 driver used throughout this study, in particular for the profiling of PolII occupancy in the somatic testis population. The yellow asterisk marks the location of the hub. **(B)** Venn diagram showing the overlap of genes bound by Zfh-1 (green), Abd-A (orange) and expressed in the soma (PolII) (blue) in *Drosophila* L3 testes, resulting in the definition of seven gene classes. Gene numbers in the respective categories are indicated. **(C)** Heat-map displaying presence of genes belonging to different signalling pathways in five major gene classes. The colour range corresponds to the fraction of genes annotated to the category that also appear in the sample: dark-red: 75% of the genes of the category are present in the sample, dark-blue: none of the genes in the category are present in the sample. **(D)** Heat-map displaying enrichment of general higher-order categories. The colour represents the p-values:dark-red < 10^−20^, dark-blue: > 0.05. Green frame highlights all genes active but not bound by Zfh-1 and Abd-A (Class 7), red frame highlights all gene classes containing active genes targeted by Zfh-1 and / or Abd-A. **(E)** Zfh-1 and Abd-A-Dam binding profiles for the *zfh-1* and *Socs36E* loci. Average peak intensity calculated from 2 repeats in dark and light green, respectively. Associated genes/transcript are shown in light orange. Testes are oriented anterior left. Scale bars, 10 μm. See also Figure S2.

Taken together, our analysis showed that the two regulatory proteins Zfh-1 and Abd- A fulfil important and distinct functions in the *Drosophila* testis stem cell system via regulation of their targets. Thus, they represent ideal candidates to identify genes controlling processes active in the early stages of the somatic support cells and critical for non-autonomously regulating the balance between stem cell self-renewal and differentiation.

### A functional gene expression atlas of the *Drosophila* testis soma

In order to identify genes active in CySCs subpopulations and their immediate progenies of the *Drosophila* larval testis, we generated datasets representing expressed genes in the *Drosophila* testis soma and germline, as well as genome-wide binding regions for the TFs Zfh-1 and Abd-A. To this end, we used Zhf-1 and Abd-A DamID (Cruciat et al., 2010; Gross et al., 2012; Tolhuis et al., 2011; van Steensel et al., 2001; Zoncu et al., 2011) and Pol II Targeted DamID (TaDa) Inoue et al., 2005; Southall et al., 2013) for mapping RNA polymerase II (Pol II) occupancy, which served as a proxy for gene activity. For TaDa, we expressed a fusion protein consisting of the *Escherichia coli* DNA adenine methyltransferase and Pol II (UAS-*Dam-PolII*) either throughout the somatic cell population using the c587-GAL4 driver (Figure 2A) or the germline lineage using the *nanos (nos)-GAL4* line (Figure S2B) (Stevens and Forgac, 1997; Van Doren et al., 1998). For conventional DamID, we generated Dam-Abd-A and Dam-Zfh-1 transgenic flies and used leaky expression from the heat-shock promoter to drive very low expression of the Dam fusion proteins (Allan et al., 2005; Tolhuis et al., 2011; van Steensel et al., 2001).

We considered the DamID strategy the best suited for our purpose for the following reasons. First, DamID had successfully been used in the past for mapping binding sites of TFs active in stem cells (Forgac, 2007; Korzelius et al., 2014). Second, low level expression of the Dam-Abd-A as well as Dam-Zfh-1 fusion proteins had no visible or molecular effect on testis development, neither in the somatic nor in the germline lineage. Fully functional adult testes formed, which showed gene expression indistinguishable from wild-type testes (Figures S2D-S2I). Third, the *c587*-GAL4 and nos-GAL4 drivers used for PolII TaDa are active throughout larval development, thus Dam-mediated DNA methylation sites represent binding events over the course of testis development. In contrast, other genomic approaches, such as RNA- or ChIP-seq, can only provide snap-shots of individual time points. After extraction from L3 testes, methylated DNA was purified and detected by hybridization to *Drosophila* Tiling Arrays. Enriched binding regions were defined by comparing PolII/Zfh-1/Abd-A methylation profiles to a Dam-alone control (see Materials and Methods). Subsequently, we called genes that had at least one enriched genomic region within 2 kb of the gene body in both DamID replicates as targets. For gene detection in TaDa experiments, we used high stringency parameters (FDR < 0.01), which led to the recovery of 5905 genes expressed in larval somatic cells (Figures 2B, S2C). This list included *zfh-1* and *abd-A* as well as many other genes known to be active in the testis soma (Table S1). For the germline, we identified 2199 expressed genes using the same strategy (Figure S2C).

Since our Dam-fusion proteins were equally expressed in the somatic and germline cell populations, whereas the endogenous Zhf-1 and Abd-A are exclusively active in the soma, we first analysed our datasets for the specificity of our targets. To this end we compared our lists of active genes identified by germline or soma TaDa with the 2686 genes that are targeted by Abd-A and/or Zfh-1. Interestingly, we found only 6% (222/2686) of the germline-restricted active genes to be associated with Abd-A and/or Zfh-1 binding events, while 94% (2464/2686) of the genes expressed exclusively in the soma were targeted by either or both TFs (Figure S2C). This result demonstrated that Zfh-1 and Abd-A interacted with cis-regulatory regions preferentially in the somatic lineage, maybe due to permissive chromatin environment or availability of suitable cofactors.

After having confirmed the specificity of our datasets, we excluded the 222 germline-specifically expressed and TF targeted genes from subsequent analyses and combined the soma-specific transcriptome with the TF binding datasets, which led us to identify 7 major classes of genes in total (Figure 2B, Table 1). First, we found somatically expressed genes bound exclusively by Zfh-1 (Class 2) or Abd-A (Class 3) or by both TFs (Class 1) (in total: 3366 genes) (Figure 2B). Based on the activity of Abd-A and Zfh-1 (Figures 1B-1D, S1A, S1B) (Deng and Lin, 1997; Leatherman and DiNardo, 2008) and the experimental set-up, we expected these classes to contain many genes expressed in CySCs and early SCCs at one point in larval testis development (and bound by Abd-A and/or Zfh-1) (Table 1). Consistently, fifteen genes known to be active, in some cases exclusively in CySCs and early SCCs were present in these three classes, including *zfh-1* (de Cuevas and Matunis, 2011; Leatherman and DiNardo, 2008), *C-terminal Binding Protein (CtBP)* (Gleixner et al., 2014; Kim et al., 2014; Leatherman and DiNardo, 2008; Petzoldt et al., 2013), *dlg1* (Murakami et al., 2004; Papagiannouli and Mechler, 2009; Sun et al., 2010), *Suppressor of cytokine signalling at 36E (Socs36E)* (Chen et al., 2013a; Colozza et al., 2011; Issigonis and Matunis, 2012), *Signal transducer and activator of transcription protein at 92E (stat92E)* (D’Angelo and Hetzer, 2008; Kiger et al., 2001), *ken and barbie (ken)* (Issigonis and Matunis, 2012; Pascual-Garcia and Capelson, 2014; Van de Vosse et al., 2013) *cubitus interruptus (ci)* (Amoyel et al., 2013; Chen et al., 2013a), *Epidermal growth factor receptor (EGFR)* (Chen et al., 2013a; Kitadate and Kobayashi, 2010), *Retinoblastoma-family protein (Rbf)* (Dominado et al., 2016; Lim and Fuller, 2012), *Ecdysone receptor (EcR)* (Ayyub et al., 2015; Dominado et al., 2016; Gonzalez et al., 2015; Li et al., 2003; 2014; Saito et al., 2009), *chickadee (chic)* (Fairchild et al., 2015; Papagiannouli et al., 2014), *punt (put)* (Hay et al., 1994; Matunis et al., 1997), *Nucleosome remodelling factor 38kD (Nurf38)* (Cherry and Matunis, 2010; Insco et al., 2009), *schnurri (shn)* (Buchon et al., 2009; Klimmeck et al., 2012; Matunis et al., 1997) and *robo2* (Southall et al., 2013; Stine et al., 2014) (Table S1). Binding profiles of selected target loci showed that strong enrichment for Zfh1-Dam or AbdA-Dam binding was found mostly around the transcription start site or within intronic regions (Figures 2E, S2J). Second, we found sets of genes targeted by both TFs (Class 4), or exclusively by Zfh-1 (Class 5) or Abd-A (Class 6), but not recovered in the PolII TaDa experiment (in total: 1663) (Figure 2B). These groups were classified as genes inactive in the testis soma and either actively repressed by Zfh-1 and/or Abd-A or neutrally bound (Table 1). And finally, a large number of genes was recovered in the soma-specific PolII TaDa experiment but not associated with Zfh-1 or Abd-A binding (2539) (Class 7) (Figure 2B). We classified these genes to be expressed throughout the soma with their expression being independent of Zfh-1 and Abd-A inputs and very likely encoding proteins that fulfil more general functions (Table 1).

**Table 1.** **Assignment of gene classes to somatic cell types in the *Drosophila* testis.** The assignment of gene classes (left panel) to gene activities in somatic cell types (CySCs, early SCCs, late SCCs) (right panel) is based on the binding profiles of Polymerase II (PolII), which indicates active genes, and of the two transcription factors Zfh-1, active in CySCs and their immediate daughter cells, and Abd-A, expressed in CySCs and early SCCs (middle panel).

| Gene Classes | Genomic Binding Profile | Assignment / Interpretation |
| --- | --- | --- |
| Class 1 | PolII ✓<br>Zfh-1 ✓<br>Abd-A ✓ | active in somatic lineage<br>(in particular in CySCs + early SCCs) |
| Class 2 | PolII ✓<br>Zfh1 ✓<br>Abd-A ✗ | active in somatic lineage<br>(in particular in CySCs + immediate daughter cells) |
| Class 3 | PolII ✓<br>Zfh-1 ✗<br>Abd-A ✓ | active in somatic lineage<br>(in particular in CySCs + early SCCs) |
| Class 4 | PolII ✗<br>Zfh-1 ✓<br>Abd-A ✓ | inactive in somatic lineage<br>(including CySCs + early SCCs) |
| Class 5 | PolII ✗<br>Zfh-1 ✓<br>Abd-A ✗ | inactive in somatic lineage<br>(including CySCs + immediate daughter cells) |
| Class 6 | PolII ✗<br>Zfh-1 ✗<br>Abd-A ✓ | inactive in somatic lineage<br>(including CySCs + early SCCs) |
| Class 7 | PolII ✓<br>Zfh-1 ✗<br>Abd-A ✗ | active in somatic lineage<br>(including late SCCs) |

### Gene functions active in the support cells of the *Drosophila* testis

To identify novel players controlling the *Drosophila* testis from the somatic side, we wanted to elucidate the processes most prominently represented in this lineage. To this end, we first determined over-represented GO terms in the different gene classes. To restrict our analysis to biologically relevant top-level categories, we carefully combined functionally related GO terms into higher-order categories, which we labelled using the major processes represented, like translation, transcriptional regulation or signalling (Table S2). Subsequently, we developed an online tool that allowed us to assess the enrichment of these higher order GO terms in a comparative fashion across multiple samples including interactive hierarchical representation (see Materials and Methods for details). Using this tool, we systematically identified biological processes that were differentially represented among the different gene classes.

In a first step, we tested our assignment of gene activities to somatic cell populations (Table 1). To this end, we compared processes overrepresented in the Zfh-1 and Abd-A active class (Classes 1, 1+2, 1+3, 1+2+3), which we assumed to be particularly required in CySCs and early SCCs (Table 1), with those enriched in the control gene class (Class 7), which we defined as essential processes in all somatic cells not controlled by Abd-A and Zfh-1 (Table 1). This analysis revealed a striking correlation of functional categories with the diverse cell types in the somatic lineage and known requirements for stem cell regulatory processes. For example, the general Class 7 exhibited a significant over-representation of the functional term “Translation”, in particular of processes associated with ribosome biogenesis and translational initiation, while this term was only weakly represented in the classes defined by Zfh-1 and Abd-A binding (Figure 2D). This finding was consistent with the low translational rates recently described for diverse stem cells, including *Drosophila* ovarian and male stem cells (Dutta et al., 2015; Slaidina and Lehmann, 2014) and mouse neural and hematopoietic stem cells (Dutta et al., 2015; Llorens-Bobadilla et al., 2015; Signer et al., 2014). Conversely, we found differentiation-related genes to be prominently overrepresented in Zfh-1 and Abd-A active classes but not in Class 7 (Figure 2D). Thus, differentiation processes seem to be primed in CySCs (and early SCCs), which is in line with functional studies in the testis (Figures 1D-1I) (Cabezas-Wallscheid et al., 2014; Leatherman and DiNardo, 2008). Interesting examples of genes within this category, which are known for controlling differentiation processes in CySCs, are the TF encoding genes *Stat92E* (Kiger et al., 2001; Southall et al., 2013), *zfh-1* and *CtBP* (Jiang et al., 2016; Leatherman and DiNardo, 2008). However, our tool recovered many other genes belonging to the “Differentiation” group so far not known for controlling the stem cell maintenance and differentiation switch in the somatic lineage of the testis, including the TFs Retinoblastoma factor (Rbf), the Myeloid leukemia factor (Mlf), Hyrax (Hyx), Female sterile (1) homeotic (Fs(1)h), and Smad on X (Smox).

Similarly, the functional terms “Signaling” and “Stimulus” were strongly enriched among the genes of the specific Zfh-1 and Abd-A active classes but not in the general soma expressed Class 7 (Figure 2D). This finding may reflect the well-known dependency of the male stem cell niche on the proper interplay of signalling pathways (Chen et al., 2013b; Kiger et al., 2001; Sinden et al., 2012). Intriguingly, components of the JAK-STAT (Cabezas-Wallscheid et al., 2014; Kiger et al., 2001; Tulina and Matunis, 2001), EGFR (Chen et al., 2013a; Kitadate and Kobayashi, 2010), Hedgehog (Amoyel et al., 2013; Michel et al., 2012), MAPK (Amoyel et al., 2016), TGFβ (Li et al., 2007) and Hippo (Amoyel et al., 2014) pathways, which had been shown to be active and functional in somatic support cells, were highly enriched in Zfh-1 and Abd-A active classes but much less in Class 7 (Figure 2C). Looking beyond the known players in the testis, we identified a number of signalling pathways so far not well studied in the *Drosophila* testis soma, including the Wnt, Toll, Juvenile hormone and Sevenless (Figure 2C). These findings now open new avenues to resolve the individual and combinatorial contribution of these signalling pathways in balancing stem cell maintenance and differentiation by controlling the hub-CySC and soma-germline communication.

Another novel feature revealed by our analysis was the high representation of immune-related processes among all active genes targeted by both TFs (Classes 1, 1+2, 1+3, 1+2+3) (Figure 2D), which is in line with findings in other invertebrate and vertebrate stem cell systems (Buchon et al., 2009; Klimmeck et al., 2012), and thus might represent a species independent core function of adult stem cells. Interestingly, we found components of the Toll and Imd pathways, in particular the Toll receptor Tollo, the NF-kappa inhibitor Cactus, the Imd downstream TF Relish (Rel), the Toll/Interleukin-1 receptor homology signalling molecule Ectoderm-expressed 4 (Ect4), the Toll downstream TF Dorsal-related immunity factor (Dif) and the Rel-activated caspase Dredd, to be among the active genes targeted by Abd-A and Zfh-1 (Classes 1, 1+2, 1+3, 1+2+3). Since these components belong to the evolutionary ancient and conserved innate immune system that had been shown recently to eliminate *Drosophila* cells perceived as unfit (Meyer et al., 2014), this could indicate that these and other immune-related genes function in stem cells as an internal cell surveillance mechanism to eliminate compromised cells, a control mechanism particularly important in this type of cell. In sum, combining genome-wide analyses of active genes with the chromatin binding behaviour of two key TFs of the *Drosophila* testis soma allowed us to generate a functionally derived high-resolution gene expression atlas of supportive cells in a complex and continuously active stem cell system.

### A Regulatory Network of Signal Processing and Signalling

In a next step, we used selected processes identified as described above and crossreferenced our gene sets with databases of physical and genetic interactions in *Drosophila* to recover functionally connected genes that either act in protein complexes or pathways.

It has been recently realized that the activity of various intracellular signalling pathways is critically dependent on vacuolar (H^+^)-ATPases (V-ATPases) (Cruciat et al., 2010; Gross et al., 2012; Zoncu et al., 2011). V-ATPases are ATP-dependent proton pumps required for acidification of subcellular compartments, membrane trafficking, pH homeostasis and protein degradation (Forgac, 2007; Inoue et al., 2005). These proton pumps are hetero-multimeric complexes composed of a cytoplasmic domain, V_1_, required for ATP hydrolysis and a transmembrane domain, V_0_, critical for proton translocation (Stevens and Forgac, 1997). We found 16 genes coding for V-ATPase subunits to be expressed in the testis soma, and selected two V_1_ subunits, Vha13 and VhaAC45 (Allan et al., 2005), for functional characterization in the *Drosophila* testis. For Vha13, we identified a uniform expression through the somatic lineage of the testis (Figures S3A, S3B), in line with our assignment of gene activities to somatic cell populations (Table 1). Consistent with their known function in acidifying vesicles for protein sorting, trafficking and turnover (Forgac, 2007), endolysosomal acidification was dramatically reduced upon interference with V-ATPase function. While Lysotracker-488, a pH sensitive green fluorescent dye, was strongly enriched in wildtype testes (Figure S3C), demonstrating the extensive formation of acidic organelles, soma-specific depletion of Vha13 resulted in a strong reduction of LysoTracker-488 signal (Figure S3D). These cellular defects had severe consequences for testis development and the somatic and germline lineages were dramatically affected. First, we observed an impairment of soma differentiation when *Vha13* or *VhaAC45* were knocked down in the soma. In these testes the CySC and early SCC marker TJ was expanded, resulting in somatic cells co-expressing TJ and the late SCC marker Eya far away from the hub (Figures S4F, S4G), which was never observed in wild-type testes (Figure S4E). The inability of somatic cells to properly differentiate also affected the germline, as spermatogonia, the transit-amplifying germ cell population, were unable to exit the mitotic cycle and over-proliferated. This resulted in the emergence of cysts containing more than 16 spermatogonial cells (Figures 3B, 3C, 3E, 3F, 3H, 3I, 3K). Concomitantly with their inability to exit mitosis, the germline also displayed differentiation defects, indicated by the presence of abnormal fusomes, germ cellspecific, spectrin-rich organelles (Deng and Lin, 1997). In wild-type testes, fusomes normally associated with GSCs and goniablasts, also termed spectrosomes, had a spherical appearance (Figure S4A), while in 2- to 16-cell spermatogonia these organelles elongated and always displayed a highly branched morphology when associated with spermatocytes (Figure S4A) (de Cuevas and Matunis, 2011). In contrast, highly branched fusomes were not detected in the germline of *Vha13* or *VhaAC45* soma-depleted testes, while spherical and dumbbell shaped fusomes associated with GSCs and gonioblasts were found at the anterior tip as well as at ectopic locations far away from the hub (Figures S4B, S4C). The observed phenotype was not due to a loss of interaction between developing germ cells and SCCs, as *Vha13* and *VhaAC45* depleted as well as wild-type SCCs labelled by F-Actin enveloped the germ cells equally well (Figures 3D, 3G, 3J). Taken together, these results showed that V-ATPases not only controlled the progression of somatic cells through the different stages of their developmental program cell-autonomously, but were also required cell non-autonomously for the completion of the transit-amplifying program and proper differentiation of germ cells into mature spermatocytes, most likely by transducing a so far unknown signal(s) to the germline.

**Figure 3:**
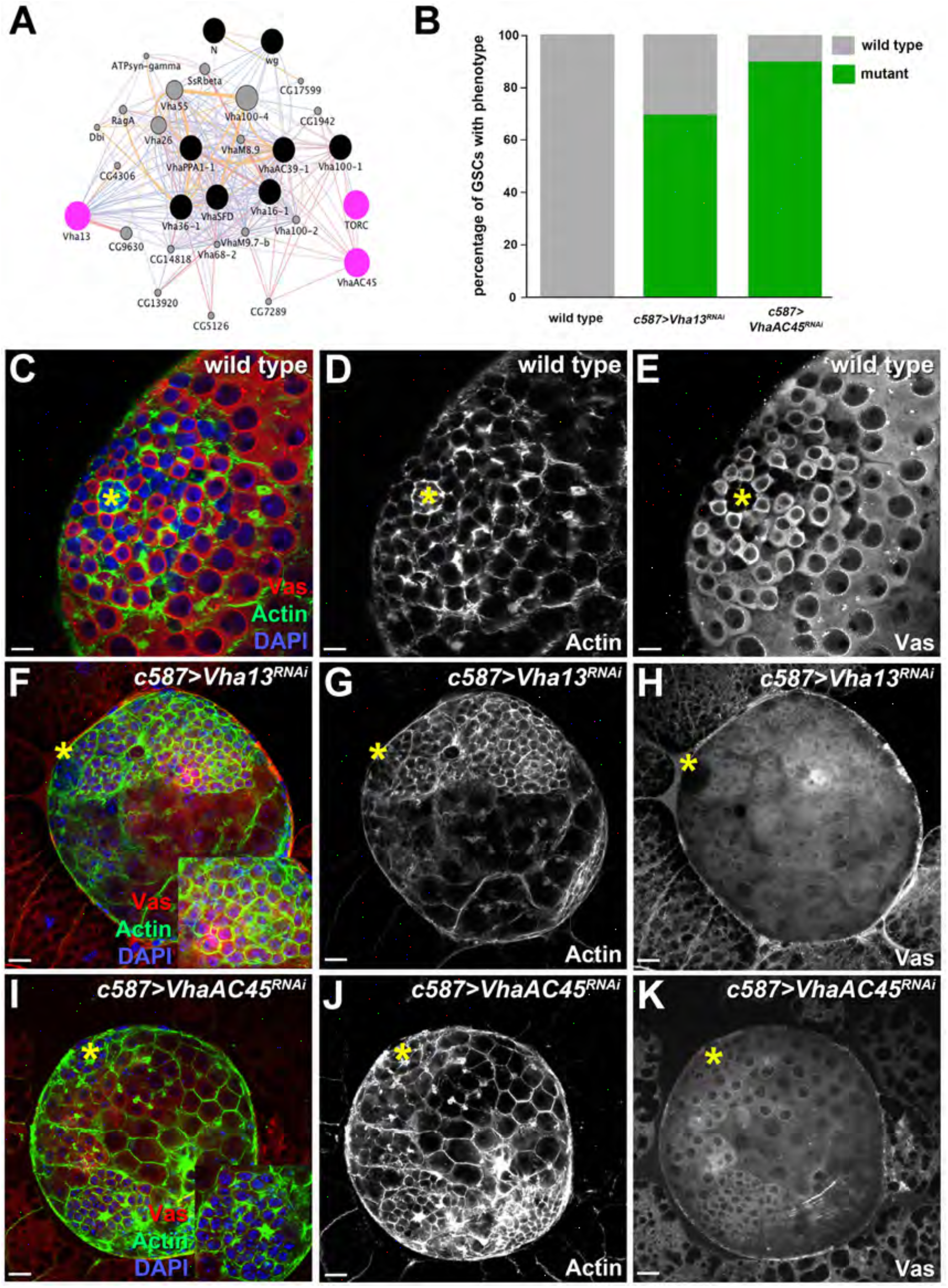
V-ATPases are required for proper development of the germline and somatic lineage in the *Drosophila* testis. **(A)** Gene network showing interactions of V-ATPase subunits and a few signalling components identified in this study, genes coloured in pink indicate those analysed in more detail. **(B)** Quantification of testes displaying GSC phenotype in wild-type, *c587>Vha13^RNAi^* and *c587*>*VhaAC45^RNAi^* animals. Green color indicates the mutant phenotype, grey color indicates the wild-type phenotype. (C-K) L3 wild-type **(C-E)**, *c587*>*Vha13^RNAi^* (F-H) and *c587>VhaAC45^RNAi^* (I-K) testes stained with Vas (red) to label the germline, Actin (green) to mark cyst cells and DAPI (blue) for the DNA. Knock-down of the V-ATPase subunits Vha13 and VhaAC45 results in over-proliferation of spermatogonia. The insets in (F, I) represent high magnification images to show the overproliferation of spermatogonia. Yellow asterisks mark the location of the hub. Testes are oriented anterior left. Scale bars, 10 μm. See also Figure S3.

Various signalling pathways, including Notch, Wnt, G-protein coupled receptors (GPCRs), receptor tyrosine kinases (RTKs) and TOR have recently been associated with functional V-ATPases (Gleixner et al., 2014; Kim et al., 2014; Petzoldt et al., 2013). Importantly, we found the TOR signaling pathway to be enriched in our datasets (Figure 2C), and network analysis using a publicly available gene- and protein- interaction database revealed an interaction of V-ATPase subunits with the TOR signalling cascade (Figure 3A). TOR signaling was implicated in the regulation of various mammalian and *Drosophila* stem cell systems (Murakami et al., 2004; Sun et al., 2010), leading us to test its role in the *Drosophila* testis. Similar to the effects observed in V-ATPase knock-down conditions, soma-specific interference with the central player of the pathway, *Tor,* resulted in an expansion of TJ positive somatic cells (Figures 4A, 4C, 4D, 4E, 4G) as well as of mitotically dividing spermatogonial cells that were unable to differentiate into mature spermatocytes (Figures 4B, 4F, 4I, 4K, S4D). However, *Tor* seemed to function also independently of V-ATPases, since we observed a strong reduction of differentiated Eya-positive SCCs in *Tor* soma-depleted testes (Figures 4H-4L), which was never the case in V-ATPase depleted testes (Figures S4E-S4G). In sum, these data suggested that V-ATPases controlled the activity of signalling pathways, including TOR, in the *Drosophila* testis and in other stem cell systems.

**Figure 4:**
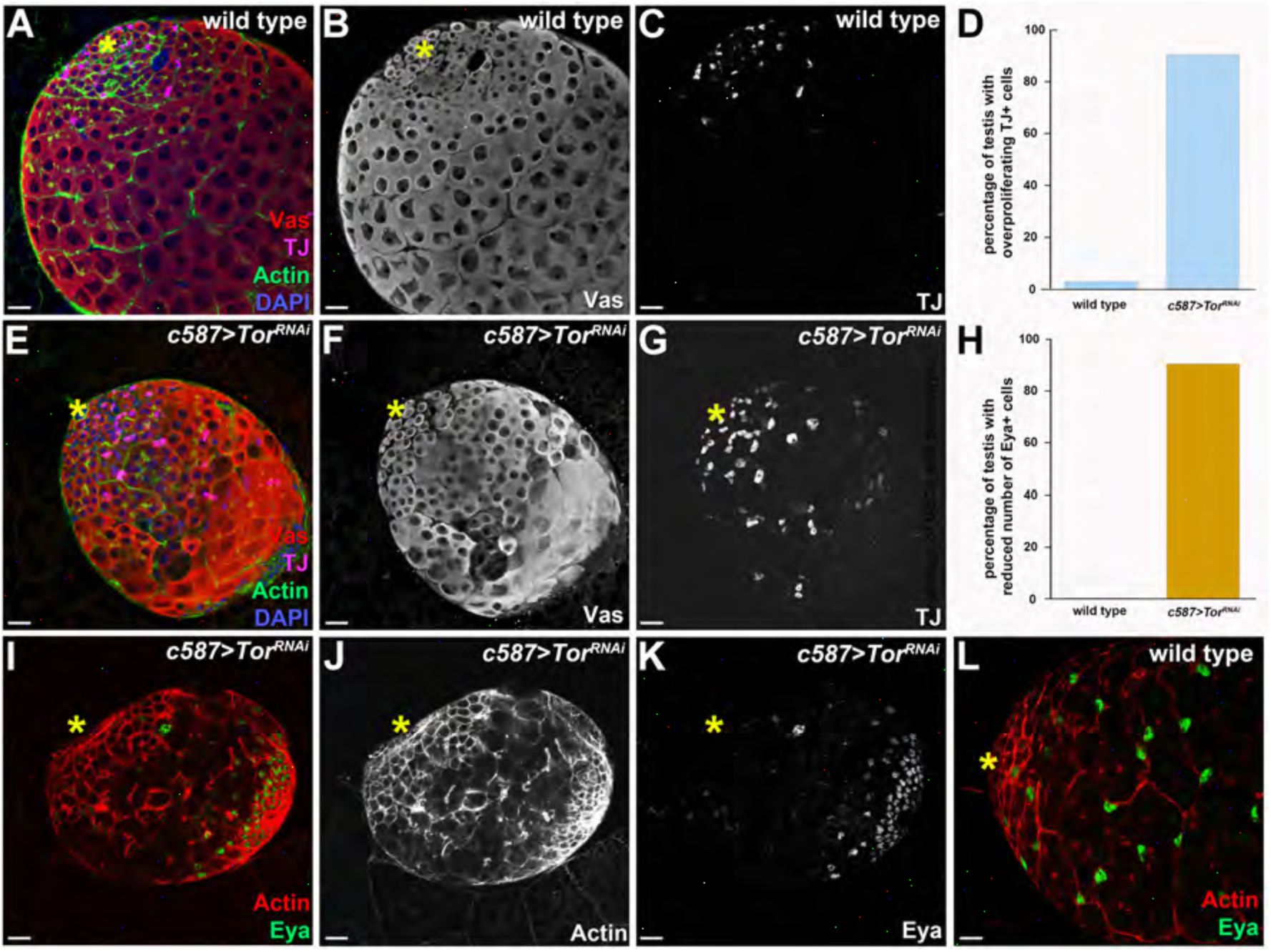
The TOR pathway is required for proper development of the germline and somatic lineage in the *Drosophila* testis. **(A-C)** L3 wild-type testes stained with Vas (red) to label the germline, TJ (magenta) to label CySCs and early SCCs, Actin (green) to mark cyst cells and DAPI (blue) for the DNA. **(D)** Quantification of testes displaying overproliferation of TJ positive cells in wild-type and *c587*>*Tor^RNAl^* animals. **(E-G)** L3 *c587*>*Tor^RNAl^* testes stained with Vas (red) to label the germline, TJ (magenta) to label CySCs and early SCCs, Actin (green) to mark cyst cells and DAPI (blue) for the DNA. (H) Quantification of testes displaying reduced numbers of Eya positive cells in wild-type and *c587*>*Tor^RNAl^* animals. **(I-L)** L3 wild-type (L) and *c587*>*Tor^RNAl^* **(I-K)** testes stained with Actin (red) to mark cyst cells and Eya (red) to label late SCCs. The number of Eya positive cells is reduced in testes with reduced TOR levels. Yellow asterisks mark the location of the hub. Testes are oriented anterior left. Scale bars, 10 μm. See also Figure S4.

### A Regulatory Network of Transport: Nuclear Pore Complex

In recent years, nucleoporins (Nups), the building blocks of the nuclear pore complex (NPC), have emerged as potential regulators of stem cell activity (Chen et al., 2013a; Colozza et al., 2011). NPCs are large nuclear envelope-embedded protein assemblies that are composed of more than 30 different Nups, creating a selective transport channel between the nucleus and the cytoplasm (D’Angelo and Hetzer, 2008) (Figure 5A). In addition to their transport-related functions, Nups have also emerged as potential regulators of chromatin organization and transcription (Pascual-Garcia and Capelson, 2014; Van de Vosse et al., 2013). Our datasets suggested that eight (out of 34) NPC-associated proteins are active in the somatic lineage of the *Drosophila* testis, three of which were associated with Abd-A or Zfh-1 binding regions. One of them was Nup153 (found in Class 1), which has been shown previously to support EGFR signalling in CySCs by promoting nuclear retention of phosphorylated Extracellular- signal Regulated Kinase (ERK), thereby influencing CySC-hub interaction and GSC development in *Drosophila* testes (Chen et al., 2013a). In order to elucidate the celltype specific function of Nups in the *Drosophila* testis, we interfered with the function of four candidates from our dataset in somatic cells by soma-specific RNAi. We tested two NPC scaffold proteins Nup44A and Nup93-1 (scaffold outer ring) and one nucleoporin located in the cytoplasmic ring, Nup358, and with Nup205, which we found to be specifically expressed in the somatic lineage and early germline (Figure 5I). In all cases, we observed a similar phenotype: an expansion of LamDm_0_ expression (Figures 5B-5F), which labels GSCs, gonialblasts and spermatogonia (Chen et al., 2013a), revealed an over-proliferation of the early germline at the expense of differentiation, as LamC-positive spermatocytes were significantly reduced in number (Figures S5A-S5E). As a consequence, 2-, 4- and 8-spermatogonia were progressively lost and the testes were filled with small LamDm_0_ labelled germ cells (Figures 5B-5F). Germline overproliferation is frequently observed when SCCs either die (Lim and Fuller, 2012) or are unable to properly encapsulate the germline (Ayyub et al., 2015; Dominado et al., 2016; Gonzalez et al., 2015; Li et al., 2003; Saito et al., 2009). Therefore, we analysed the distribution of βPS-integrin, the β-chain of the integrin heterodimer decorating the membranes of CySCs and SCCs (Papagiannouli et al., 2014) as encapsulation marker. We found that in the absence of Nup205 SCCs were indeed unable to properly wrap the over-proliferating germ cells (Figures 5J, 5K). Furthermore, we observed a progressive loss of TJ-positive cells in *c587::nup205^RNA^* testes (Figure 5H) in comparison to wild-type testes (Figure 5G), which might explain the cell nonautonomous effect on the germline. Consistent with a potential loss of these cells by cell death, the viability of the somatic cell population was restored when apoptosis was prevented in *c587::nup205^RNA^* larval testes by co-expression of the anti-apoptotic baculovirus protein p35 (Figures 5F, 5L-5O) (Hay et al., 1994). Strikingly, suppression of apoptosis seemed to partially rescue the developmental defects, since *c587::p35;nup205^RNA^* testes contained normal spermatocytes and relatively normal spermatogonial cysts reappeared (Figures 5L, 5O). These results indicated that the germline lineage was at least partially able to enter the transit-amplifying (TA) program, the early step of germline differentiation in which GSC daughters execute four rounds of mitotic divisions before entering terminal differentiation (Insco et al., 2009), even in the absence of Nup205 when the somatic cell population was kept alive. Importantly, the restoration of TJ-positive CySCs and early SCCs (Figures 5M, 5N), the reappearance of spermatogonial cysts shown by Actin staining (Figures 5L, 5M) and the suppression of germ cell over-proliferation evidenced by LamDm_0_ staining (Figures 5M, 5N) indicated that the processing of factors in the somatic lineage, which ensure germline differentiation, is at least to some extent re-established.

**Figure 5.**
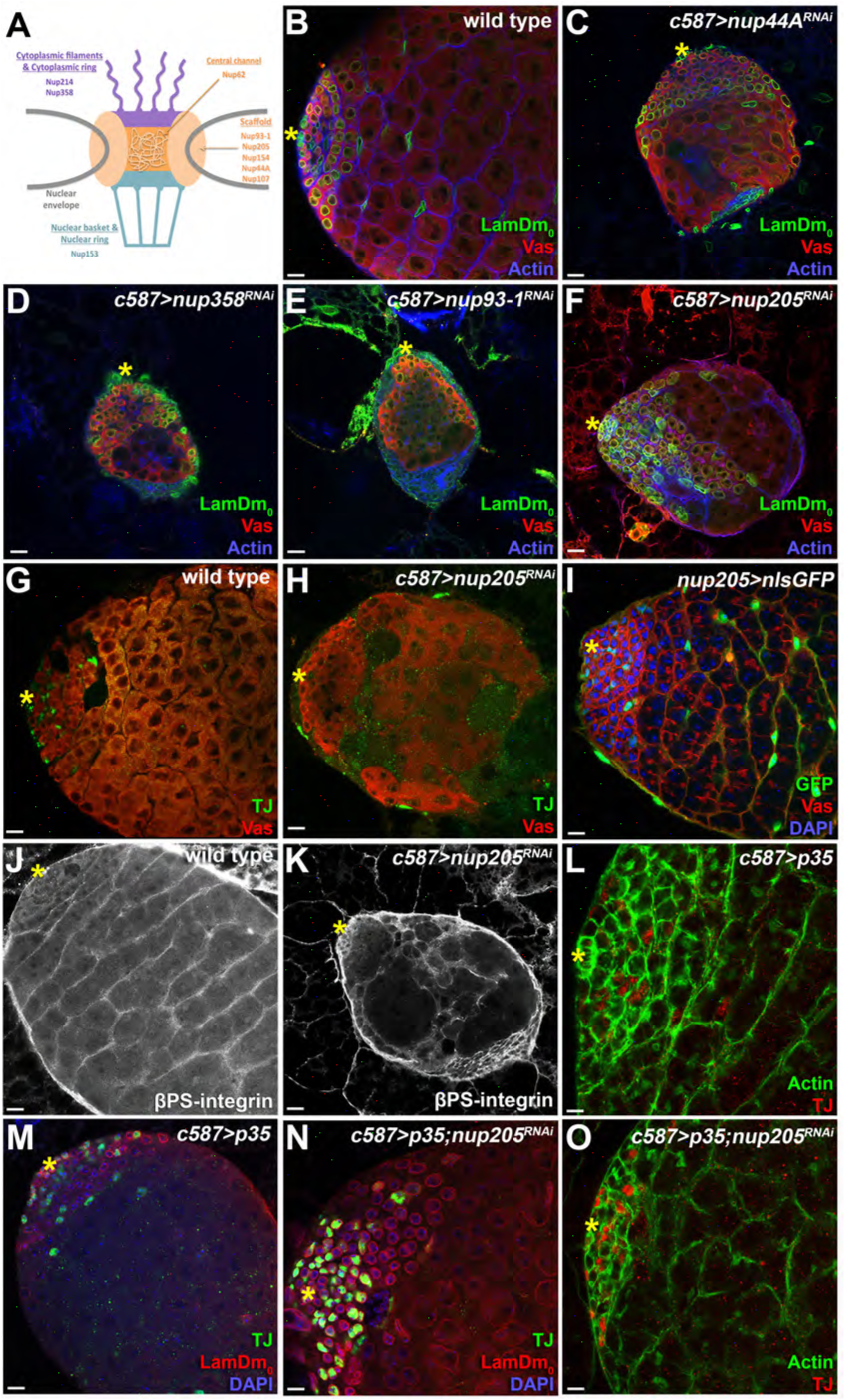
Nucleoporins have a critical function within the cyst cell population and cell non-autonomously control the germline of the *Drosophila* testis. **(A)** Schematic diagram of the nuclear pore structure depicting the localization of the nucleoporin proteins recovered in this study. **(B-F)** L3 wild-type (B), *c587*>*nup44A^RNAi^* (C), *c587*>*nup358^RNAi^* (D), *c587*>*nup93-1^RNAi^* (E) and *c587*>*nup205^RNAi^* (F), testes stained for Vas (red) to mark the germline, Actin (blue) to label cyst cells and germline fusomes, and LamDm_0_ (green) to indicate early germ cells (GSCs, gonialblasts and spermatogonia). Knock-down of nucleoporins results in over-proliferation of early germ cells. **(G, H)** L3 wild-type (G) and *c587*>*nup205^RNAi^* (H) testes stained for Vas (red) to label the germline and TJ to label CySCs and early SCCs. (I) *nup205*>*nlsGFP* L3 testis showing that Nup205 is exclusively expressed throughout the somatic lineage. **(J, K)** L3 wild-type (J) and *c587*>*nup205^RNAi^* (K) testes stained for βPS-integrin (white) to indicate cyst cells. (L) L3 *c587>p35* control testis stained for TJ (red) to label CySCs and early SCCs and Actin (green) to label cyst cells and germline fusomes. **(M, N)** L3 *c587>p35* (M) and *c587*>*p35;nup205^RNAi^* (N) testes stained for TJ (green) to label CySCs and early SCCs, LamDm_0_ (red) to indicate early germ cells (GSCs, gonialblasts and spermatogonia and spermatogonia) and DAPI (blue) for the DNA. The analysis shows that inhibition of apoptosis restores the somatic lineage, allowing the germline to enter the transit-amplifying program.**(O)** L3 *c587*>*p35;nup205^RNAi^* testis stained for TJ (red) to label CySCs and early SCCs and Actin (green) to label cyst cells and germline fusomes. Testes are oriented anterior left. Scale bars, 10 μm. See also Figure S5

Taken together, these results showed that Nups were critically required in SCCs enabling them to properly encapsulate the neighbouring germ cells, thereby restricting proliferation of spermatogonia and promoting their differentiation into spermatocytes. It will be of high relevance for our understanding of stem cell systems in the future to unravel whether Nups ensure the survival, differentiation or adhesiveness of somatic cells by selectively controlling nuclear transport, chromatin organization or transcriptional control.

### A Comparative Analysis of *Drosophila* and Mammalian Stem Cell Systems

Mining our dataset with our analysis tool allowed us to identify major overrepresented and functionally relevant processes in the soma of the *Drosophila* male stem cell system (Figure 2D). Strikingly, some of the identified genomic signatures have also emerged in other stem cell systems, like the activation of immune-related genes (Buchon et al., 2009; Klimmeck et al., 2012). Therefore, we hypothesized that conserved genomic programs may exist to control the behaviour of adult stem cells irrespective of species or niche type. Conversely, system-specific regulators would allow different stem cell types to respond to and interact with their typical yet diverse microenvironments. To test this hypothesis, we extended our online analysis tool to allow the inclusion of published expression data. To compare divergent adult stem cell systems within *Drosophila,* we added datasets on neural stem cells, the so-called neuroblasts (NBs) (Southall et al., 2013) and intestinal stem cells (ISCs) (Dutta et al., 2015). Furthermore, to delineate common and divergent processes across functionally, but also evolutionary diverse stem cell systems between species, we included data from mammalian hematopoietic stem cells (HSCs) into our analysis. These datasets were generated using diverse technologies, cell sorting and RNA-seq for ISCs (Dutta et al., 2015) and HSCs (Cabezas-Wallscheid et al., 2014) while NBs (Southall et al., 2013) and the testis soma were profiled using DamID. For NBs all genes with a PolII DamID peak at the transcriptional start site were considered as expressed, for the testis soma we used the Abd-A and Zfh-1 active genes (Class 1+2+3), while for ISCs and HSCs the top 50% of detected transcripts were analyzed. Our comparative analysis revealed a number of striking correlations. First of all, we identified a small number of processes to be similarly represented in all systems, in particular cell growth and cell cycle, which were significantly enriched in all four stem cell systems (Figure 6C). Interestingly, ISCs had a slightly higher enrichment of both processes, which is in line with the rapid cellular turnover of the gut (Jiang et al., 2016). Furthermore, we found that processes related to cytoskeleton and cell adhesion were moderately over-represented (Figure 6C), which is in line with intimate stem cell - niche association, critical for long-term maintenance and function of stem cell systems (Chen et al., 2013b). In addition, several signatures were found to be different in the various stem cell systems, including a high representation of stem cell-related processes in neuroblasts, of metabolic processes in HSCs and of the immune response in the testis soma (Figure 6C). Using a comparative analysis makes these findings highly relevant, as they could represent system-specific adaptations and thus be crucial for the activity and function of individual stem cell systems.

**Figure 6.**
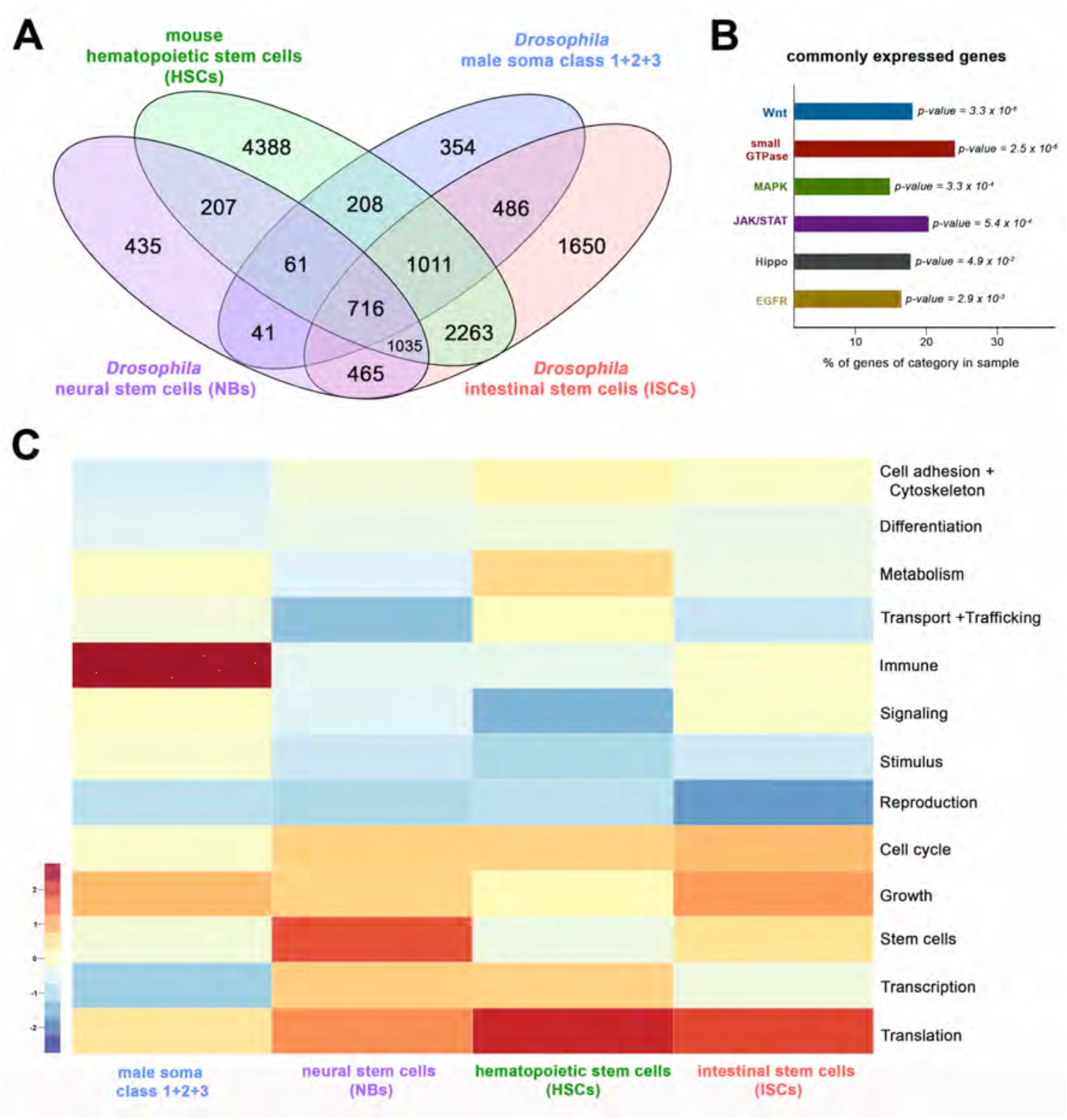
Comparative analysis of different stem cell systems. **(A)** Comparison of the transcriptome of *Drosophila* neural stem cells (purple), *Drosophila* intestinal stem cells (ISCs) (red), *Drosophila* male somatic stem cells (CySCs) and early SCCs (blue) as well as mouse hematopoietic stem cells (green). 716 genes were found in all four conditions, 354 genes are specific in testis soma dataset, 1650 in *Drosophila* intestinal stem cells, 435 in *Drosophila* neural stem cells and 4388 in mouse hematopoietic stem cells. **(B)** Enrichment of signaling pathways in the 716 commonly expressed genes from all four stem cell systems. The significance of the enrichment is presented as p-value. The length of the bars represents the fraction of genes annotated to the pathway within the sample. The p-value was calculated using Fisher test. **(C)** Heat-map displaying presence of genes belonging to higherorder categories in different stem cell systems. The colour range corresponds to the centered and scaled (per column) fraction of genes annotated to the category that also appear in the sample: red colour represents high values, blue colours low fractions of genes in the category, which are also present in the sample. Rows and columns are hierarchically clustered using Euclidean distance with complete linkage.

Building on the finding of shared and divergent functional categories, we next searched for individual genes common to all datasets. In order to do so, we used orthology mapping to convert mouse gene identifiers into *Drosophila* gene identifiers. Intersecting whole transcriptome profiling of FACS sorted mouse HSCs (Cabezas-Wallscheid et al., 2014) with the *Drosophila* datasets resulted in the identification of 716 transcripts common to all systems (Figure 6A). Since it is known that extensive communication between cells is crucial in all stem cell centers, we asked which signaling pathways were enriched among these commonly expressed genes. Using our analysis tool, we found the Wnt, MAPK JAK/STAT, EGFR and the Hippo pathways, which are all known for their functions in diverse stem cells, to be significantly overrepresented in all four stem cell system (Figure 6B). Since our study had shown that V-ATPases and nucleoporins are critical for activity and integrity of the *Drosophila* male stem cell system (Figures 3, 4, 5), we asked whether these complexes may also be of relevance across stem cell systems. We found two V-ATPase subunits, Vha44 and Vha16-1, and three nucleoporins, Nup44A, Nup107 and Nup153, to be represented in all four datasets, including the mammalian HSCs. Strikingly, Nup153 has been shown to play a critical role in somatic stem cells of the *Drosophila* testis (Chen et al., 2013a) and just recently in mouse embryonic stem cells for the maintenance of their pluripotency (Jacinto et al., 2015), while Vha44 had been implicated in proliferation control (Petzoldt et al., 2013). To test whether their functional relevance extends to an unrelated adult stem cell system, we interfered with the function of Nup44A and Nup153 in *Drosophila* ISCs. RNAi using the *escargot (esg)-* GAL4 driver (Manseau et al., 1997; Micchelli and Perrimon, 2006), resulted in a complete loss of all ISCs for Nup44A (Figures S7A-S7C), while interference with Nup153 had no effect in the intestine (Figures S7A, S7D) despite its prominent role in the testis (Chen et al., 2013a). Strikingly, knock-down of Vha16-1 in the *Drosophila* testis resulted in an over-proliferation of TJ positive CySCs and early SCCs (Figures S6A, S6C, S6J, S6L), while a dramatic stem cell loss was observed in ISCs (Figures S7A, S7C). These results supported the notion that genes commonly expressed across stem cell systems have a high probability to be functionally relevant, but also demonstrated that regulatory activity of these genes may be system specific.

To test this more systematically, we focused on transcription factors, since many TFs had been demonstrated to critically control the balance between self-renewal and differentiation in diverse stem cell systems (Takahashi and Yamanaka, 2006; Yamanaka and Blau, 2010). Intersecting all datasets, we identified a set of 21 proteins with annotated TF activity to be commonly expressed in all four stem cell systems (Table 2, Table S3), many of them being associated with the higher-order GO term “Differentiation”. This set included CtBP, which is important for stem cell maintenance in the *Drosophila* testis (Leatherman and DiNardo, 2008) and for integration of environmental signals to regulate neural stem cell state in chick embryos (Dias et al., 2014) and the ETS domain containing transcription factor Pnt-P1, known to promote the generation of intermediate neural progenitors in *Drosophila* larval brains (Dias et al., 2014) and the ETS domain containing transcription factor Pnt-P1, which is known to promote the generation of intermediate neural progenitors in *Drosophila* larval brains (Zhu et al., 2011). Another well studied regulator in this set was Rbf, required for cell cycle exit and differentiation of the somatic and germline stem cells in *Drosophila* testis (Dominado et al., 2016). Interestingly, the mammalian Retinoblastoma-like protein 1 (RBL1) controls HSC quiescence and the balance between lymphoid and myeloid cell fates in the hematopoietic system (Viatour et al., 2008). To this test the hypothesis that commonly expressed genes have a high functional relevance, we reduced the activity of 18 of the 21 TFs in two different stem cell types. Using the *c587*-GAL4 and *esg*-GAL4 drivers for RNAi-based interference, we analyzed gene function in somatic cells of the *Drosophila* testis and in ISCs of the *Drosophila* intestine, respectively. Consistent with our hypothesis, we found 12 (67%) of the commonly expressed TFs to critically control stem cell maintenance and proliferation. Five of them, Mlf, CtBP, Hyx, Fs(1)h, and Smox, affected stem cell control in both systems, while three TFs, Mnt, Diabetes and obesity regulated (DOR) and Rbf, caused defects specifically in the *Drosophila* testis and four TFs, Armadillo (Arm), Checkpoint suppressor 1-like (CHES-1-like), Ecdysone-induced protein 75B (Eip75B) and Nuclear factor Y-boxB B (Nf-YB) in the *Drosophila* intestine (Table 2, Table S3). Interestingly, three TFs seemed to have a similar function in both stem cell systems, as knock-down of Hyx, a member of the Paf1 protein complex controlling a variety of transcriptional processes (Mosimann et al., 2009), CtBP and the Bromodomain protein Fs(1)h resulted in a reduction of TJ positive somatic cells in the testis (Figures 7A, 7C, 7G, 7I, 7J, 7L) and of *Delta-lacZ* labeled ISCs in the intestine (Figures 7M-7T). Defects in somatic cell differentiation of the testis affected also the ability of cyst cells to support the germline resulting in reduction of spermatogonial cysts and premature differentiation (Figures 7B, 7H, 7K). This suggested that these TFs control germline development in the testis cell nonautonomously. A similar decrease in stem cell number was observed when the activities of Eip74EF, Smox, Arm and CHES-1-like were knocked-down in the *Drosophila* intestine (Figures S7E-S7J), while these TFs seemed to have no critical function in the testis soma, as a reduction of their activity had no effect on the somatic or germline lineage (Table 2, Table S3). In contrast, DOR, a transcriptional coactivator of Ecdysone receptor signalling (Francis et al., 2010), and Rbf seemed to be required only in the *Drosophila* testis, as interference with their activities increased the number of TJ positive somatic cells (Figures S6A, S6C, S6D, S6F, S6G, S6I) and induced overproliferation of early germline cells in *Rbf^RNA^* (Figures S6B, S6E) and germline differentiation defects in *DOR^RNAi^* testes (Figures S6B, S6H) (Dominado et al., 2016), but produced no phenotypic changes in the intestine (Table 2, Table S3, Figures S7E, S7L). Only one of the 18 TFs tested, Nuclear factor Y-box B (Nf-YB), seemed to negatively control stem cell proliferation in the *Drosophila* intestine, as the number of *esg*>*GFP* positive cells was increased in Nf-YB depleted midguts (Figures S7E, S7K). Interference with six TFs had no consequence in either stem cell system (Table 2, Table S3), which could be either due to the presence of redundant functions or the activity of these TFs being not essential for stem cell maintenance and proliferation or the inability to sufficiently reduce TF concentrations in the stem cell populations.

**Figure 7.**
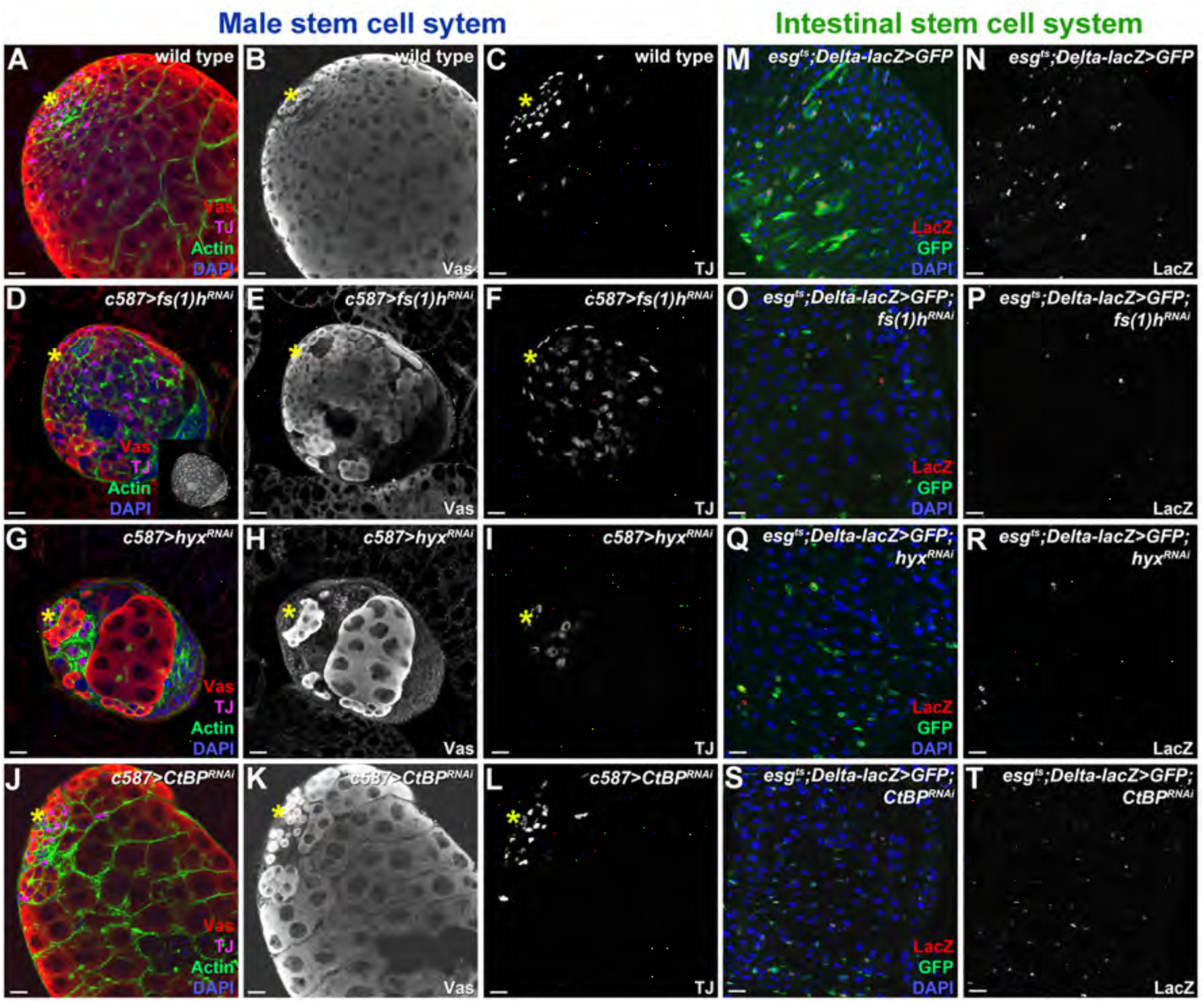
Functional analysis of commonly expressed TF encoding genes in two *Drosophila* stem cell systems. **(A-L)** L3 wild-type (A-C), *c587*>*fs(1)h^RNAi^* (D-F), *c587*>*hyx^RNAi^* (G-I), *c587*>*CtBP^RNAi^* (J-L) testes stained with Vas (red) to label the germline, TJ (magenta) to label CySCs and early SCCs, Actin (green) to mark cyst cells and DAPI (blue) for the DNA. **(M-T)** *esg^ts^*;*Delta-lacZ*>*GFP* control (M, N), *esg^ts^*;*Delta-lacZ*>*GFP;fs(1)h^RNAi^* (O, P), *esg^ts^*;*Delta-lacZ*>*GFP;hyx^RNAi^* (Q, R) and *esg^ts^*;*Delta-lacZ*>*GFP;CtBP^RNAl^* (S, T) adult guts stained for LacZ (red) to label the stem cells, GFP (green) to mark intestinal stem cells and their progenies, the enteroblasts, and DAPI (blue) for the DNA. Depletion of *fs(1)h* results in over proliferation of early germ cells indicated by bright DAPI staining (inset in d) and TJ positive cells in the testis (F), while adult midguts with reduced *fs(1)h* levels show an almost complete loss of intestinal stem cells (ISCs) (P). Interference with Hyx activity in the testis results in significant reduction of somatic cells (I) and premature differentiation of the germline (H), in adult midguts the number of ISCs is reduced (R). Knockdown of *CtBP* in the testis leads to reduction of the somatic cell population (L) and fewer spermatogonial cysts (K), in adult midguts the number of ISCs is reduced (T). Yellow asterisks mark the location of the hub. Testes are oriented anterior left. Scale bars, 10 μm. See also Figures S6 and S7.

**Table 2.**
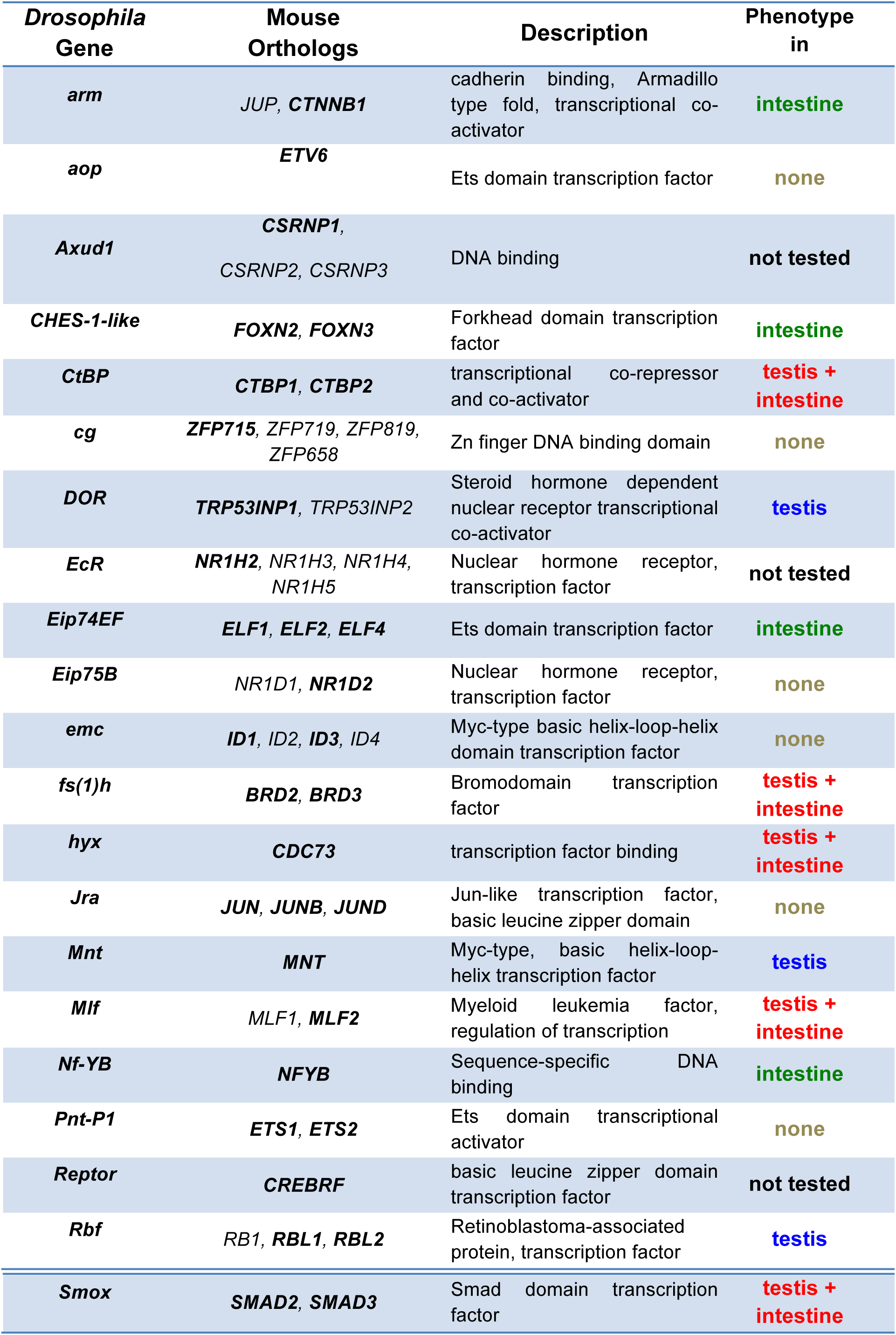
**Genes coding for transcription factors identified as commonly expressed in *Drosophila* male stem cells and progenies, intestinal stem cells and neural stem cells as well as in mouse hematopoietic stem cells.** The first column summarizes all transcription factor-encoding genes identified in all four stem cell systems. In the second column, all mouse orthologs of the *Drosophila* genes shown in the left column are presented, those in bold are the ones expressed in hematopoietic stem cells according to Cabezas-Wallscheid et al.(Cabezas-Wallscheid et al., 2014). The third column displays the description of the individual genes. The fourth column shows the results of the RNAi based interference study in *Drosophila* larval testes and adult midguts.

In sum, combining expression data from diverse stem cell system for comparative analysis using our online tool resulted in the definition of a unique core gene set. This core gene set contains an unusually high proportion of functionally relevant stem cell regulators. At 67% the phenotypic rate of this core gene set was 5 to 10 fold higher compared to phenotypes found in stem cells by genome-wide RNAi screening (Neumüller et al., 2011; Zeng et al., 2015). Importantly, the fact that loss-of-function of many of the commonly expressed genes caused divergent phenotypes in the different stem cell systems highlighted that these genes are not simply required for general cell survival, but rather that they fulfill specific regulatory roles across systems. Thus, leveraging the power of comparative transcriptomics, our datasets and online tool provide a powerful resource to quickly identify processes or genes with important functions in diverse stem cell types.

## DISCUSSION

Using cell type-specific transcriptome profiling and *in vivo* transcription factor binding site mapping together with a dedicated interactive data analysis tool, we comprehensively identified genes that are potentially involved in controlling proliferation and differentiation within a stem cell support system. Importantly, many candidates we have functionally tested not only were required within the soma, but also had non cell-autonomous functions in the adjacent (germline) stem cell lineage.

We have identified an interconnected network of transcriptional regulators, which play an important role in maintenance and differentiation of both germline and somatic cell populations, as well as novel signalling pathways, including the TOR cascade and signal processing proton pumps, the V-ATPases, as novel regulators in the *Drosophila* male stem cell system.

V-ATPases in particular have been implicated in regulation of various cellular processes, not only in invertebrates but also in vertebrates. For example, the V-ATPase subunit V1e1 had been shown previously to be essential for the maintenance of neural stem cells in the developing mouse cortex, as loss of this subunit caused a reduction of endogenous Notch signaling and a depletion of neural stem cells by promoting their differentiation into neurons (Lange et al., 2011). Furthermore, two independent studies revealed that V-ATPase subunits and their isoforms were required for proper spermatogenesis in mice, in particular for acrosome acidification and sperm maturation (Imai-Senga et al., 2002; Jaiswal et al., 2014). Thus, it is tempting to speculate that these proton pumps also have important functions in the stem cell pool of the mammalian testis and very likely many other stem cell systems, and we provide some evidence for their crucial role in ISCs.

Besides signal processing proton pumps, this work also uncovered nuclear transport associated proteins, the nucleoporins, as important control hubs in the somatic lineage of the *Drosophila* testis. This is of particular interest, since cell-type specific functions of Nups have been identified only recently and may represent a critical feature of different stem cell systems. Examples include that Nup96 heterozygous mice show defects in the innate and adaptive immune response, whereas Nup358 heterozygous mice exhibit altered glucose metabolism in the central nervous system and change into electrophysiological responses of photoreceptor neurons (Aslanukov et al., 2006; Cho et al., 2009; Raices and D’Angelo, 2012); Nup210 has been shown to function in myogenic and neuronal differentiation (D’Angelo and Hetzer, 2008; Olsson et al., 2004),while Nup133 is involved in neural stem and progenitor cell differentiation (Lupu et al., 2008); and finally some Nups are even implicated in mitotic division, like Nup358 which plays a role at kinetochores and Nup98 which regulates the anaphase promoting complex (APC) and mitotic microtubule dynamics to promote spindle assembly (Salsi et al., 2014). Interestingly, it has been shown just recently that Nups play a critical role in regulating the cell fate during early *Drosophila* embryogenesis thereby contributing to the commitment of pluripotent somatic nuclei into distinct lineages (Hampoelz et al., 2016) and our results now suggest that they may play a similar role in controlling the transition of continuously active adult stem cells towards differentiation. The next challenge will be to unravel how variations in the composition of an essential and basic protein complex like the NPC causes differential responses of cells, in particular in stem cells and their progenies.

Our new datasets in conjunction with the versatile and easy-to-use analysis tool allowed us to identify a substantial number of novel stem cell regulators for detailed mechanistic characterization. Importantly, our analyses have shed first light on processes and genes shared between diverse invertebrate and vertebrate stem cell systems, as well as uncovered functionally relevant differences. Owing to its flexibility and the option to include datasets from any species, our online tool represents a valuable resource for the entire stem cell community. It not only provides an open platform for data analysis, but leverages the power of comparative analysis to enable researchers mining genomic datasets from diverse origins in an meaningful and intuitive fashion.

## METHODS

### Fly stocks and husbandry

*Oregon R* was used as a wild type stock. The following stocks were obtained from the Bloomington Stock Center (BL) Indiana or the Vienna Drosophila RNAi Center: *UAS-abdA-RNAi^TRiP.JF03167^*, *UAS-abdA-RNAi^v106155^*, *UAS-arm-RNAi^v107344^*, *UAS-aop-RNAi^35404BL^*, *UAS-nup93-RNAi^TRiP.HMS00850^*, *UAS-nup93-1-RNAi^TRiPH.MS00898^ UAS-nup358-RNAi^TRiP.HMS00865^*, *UAS-nup358-RNAi^TRiP.HMS00803^*, *UAS-nup44A-RNAi^TRiP.HMS01825^*, *UAS-mCD8-GFP^5139BL^*, *UAS-abdA* (Michael Akam), *UAS-zfh-1*, *c833-GAL4*, *nanos-GAL4 (nanos-Gal4VP16)*, *UAS-Vha13-RNAi^38233BL^*, *UAS-Vhaac45-RNAi^38522BL^*, *UAS-Vha16-1 RNAi^v104490^*, *UAS-Vha44 RNAi^v101527^*, *UAS-CtBP-RNAi^32889BL^*, *UAS-CtBP-RNAi^v107313^*, *UAS-CHES-1-ike^26760BL^*, *UAS-CHES-1-ike^v105641^*, *UAS-Eip74EF-RNAi^v105301^*, *UAS-fs(1)h-RNAi^35139BL^*, *UAS-fs(1)h-RNAi^v108662^*, *UAS-hyx-RNAi^31722BL^*, *UAS-hyx-RNAi^31722BL^*, *UAS-Rbf-RNAi^65929BL^*, *UAS-pnt-RNAi^35038BL^*, *UAS-mnt-RNAi^29329BL^*, *UAS-mnt-RNAi^v101991^*, *UAS-Mlf-RNAi^41820BL^*, *UAS-Mlf-RNAi^v100719^*, *UAS-Nf-YB-RNAi^v103655^*, *UAS-Nf-YB-RNAi^57254BL^*, *UAS-Jra-RNAi^31595BL^*, *UAS-cg-RNAi^34668BL^*, *UAS-DOR-RNAi^60389BL^*, *UAS-Eip75B-RNAi^43231BL^*, *UAS-emc-RNAi^24768BL^*, *UAS-emc-RNAi^v100587^*, *UAS-Smox-RNAi^v105687^*, *UAS-TOR-RNAi^33951^*, *UAS-TOR-RNAi^34639BL^*, *UAS-nup205-RNAi^v38608^*, *UAS-nup205-RNAi^v38610^*, *UAS-nup44A-RNAi^v106489^*, *Vha 13-GFP^50828^*, *P{GawB}Nup205179Y*, *c833-GAL4* (drives expression in hub, CySCs and SCCs) (Papagiannouli and Mechler, 2009), *nanos-GAL4-VP16* (Van Doren et al., 1998), *c587-GAL4* (drives expression in CySCs and all SCCs) and *c587-GAL4*; *tub-Gal80^ts^* were provided by Margaret Fuller, the *UAS-nlsGFP* and *Delta-lacZ* lines were provided by Bruce Edgar. Other fly stocks used in this study are described in FlyBase (www.flybase.org). *pUAST-NDam-Myc* control flies for performing the DamID (Choksi et al., 2006) and *UAS-LT3-NDam*, *UAS-LT3-NDam-PolII* for performing TaDa (cell-type specific Targeted DamID) (Southall et al., 2013) in L3 testes were provided by Andrea Brand. All *UAS-gene^RNAl^* stocks are referred to in the text as *gene^RNAl^* for simplicity reasons. Knockdowns were performed using the *UAS-GAL4* system (Brand and Perrimon, 1993) by combining the *UAS-RNAi* fly lines with cell-type specific GAL4 drivers described above (29°C). Transgenes were expressed in ISC/EBs under the temperature sensitive control of the *esg-GAL4*, *tub-GAL80ts*, *UAS-GFP*; (*esg^ts^*) expression system as described in previous study (Micchelli and Perrimon, 2006). For all RNAi experiments in Drosophila intestine, flies were collected within 1-3 days after eclosion and were maintained at 29°C for 10 days to induce the RNAi effect.

### Immunofluorescence staining and microscopy

Whole mount testes immunostaining testes were dissected in PBS, fixed for 30min in 8% formaldehyde (PFA), rinsed in 1% PBX (1% Triton-100x in PBS) and blocked in 5% Bovine Serum Albumin in 1% PBX. Testes were incubated with primary antibodies over-night at 4^o^C and the following day with the secondary antibodies for 2h at room temperature in the dark (Papagiannouli et al., 2014). For testes immunostaining in the presence of GFP, 1% PBT (1% Tween-20 in PBS) was used instead of 1% PBX in all steps. Testes were mounted in ProLong^®^ Gold Antifade (Thermo Fischer Scientific) or VECTASHIELD (Vector Laboratories). For gut staining, tissues were dissected and fixed for 30 min in 4% PFA and washed in PBS, then blocked in 1% BSA for 30 min. The samples were incubated overnight at 4^o^C with the primary antibody (mouse anti-beta-galactosidase 1:1500, DSHB), washed 4 times with PBS (15 min each) and then incubated with secondary antibodies for 2 h at room temperature. After 4 times wahing with PBS, the tissues were mounted in Vectashield (Vector Laboratories, USA) and imaged using a Leica SP5 confocal microscope(Zhou et al., 2017). The monoclonal antibodies used in this study: anti-Armadillo N7A1 (1/10; mouse), anti-FasIII (1/100), anti-eya10H6 (1/100; mouse), anti-α-spectrin-3A9 (1/100; mouse), anti-Vasa (1/10; rat), anti-βPS-integrin-CF.6G11 (1/10; mouse), anti-LamC-LC28.26 (1/10; mouse), anti-LamDm_0_-ADL101 (1/10; mouse), were obtained from the Developmental Studies Hybridoma Bank developed under the auspices of the NICHD and maintained by The University of Iowa, Department of Biological Sciences, Iowa City, IA 52242. Polyclonal anti-Vasa-dC13 (goat; 1/20) and anti-abdA-dH17 (goat; 1/20) were from Santa Cruz Biotechnology. Monoclonal anti-Phosphotyrosine Antibody clone 4G10 (mouse; 1/200) was from Millipore, guinea-pig anti-TJ (1:5000) and rabbit anti-Zfh1 (1:2000) polyclonal antibodies were gifts of Dorothea Godt and Ruth Lehmann, respectively. Filamentous Actin (F-actin) was stained with Alexa Fluor phalloidin 488, 546 or 647 (1/300, Thermo Fischer Scientific) and DNA with DAPI (Thermo Fischer Scientific). Following secondary antibodies were used: Cy5-conjugated goat anti-mouse IgG, Cy3- conjugated goat anti-mouse IgG and Cy3-conjugated goat anti-rat IgG (Jackson Immunochemistry, PA). For the Lysotracker assay, dissected 3^rd^ instar larval testes were incubated in M3 medium containing LysoTracker^®^ Green DND-26 (Molecular probes) for 10 min. After washing in M3 medium the testes were quickly subjected to imaging.

Confocal images were obtained using a Leica system SP8 (1024x1024pix, 184μm image frame) (COS, University of Heidelberg). Pictures were finally processed with Adobe Photoshop 7.0.

### DamID (DNA adenine methyltransferase identification)

Analysis was performed as previously reported (Papagiannouli et al., 2014). For a detailed protocol see “Supplementary Materials and Methods”.

### DamID analysis

Logarithmic intensities were quantile normalized and averaged over experiments using R (http://www.r-project.org/) package Starr (version 1.28.0) (Zacher et al., 2010). Due to intrinsic biological differences between experiments, two approaches were used to generate gene lists. For Abd-A and Zfh-1, peaks were called using the script “find_peaks” (Marshall and Brand, 2015) with parameters quantile >= 0.95 and false discovery rate (FDR) <= 0.01. To find an associated gene the script “peaks2gene” (from the same publication) was used with parameter gene_pad=2000 and *Drosophila* genome annotation release 5.1. To calculate PolII occupancy and associated FDR the script “polii.gene.call” (Southall et al., 2013) with FDR<=0.01 was used.

### GO term enrichment analysis

A custom set of high-order GO terms was created by binning annotated genes into broad categories. First, we chose a set of keywords that represented different biological processes. The generic GO slim set taken from the GO consortium was then filtered and sorted into the categories by these keywords. This was followed by manually reviewing the categories to add terms or remove them when necessary, always with the aim to achieve a comparable depth of information in each category. The categories are shown in the table (Table S2). The category set for signaling pathways was created in a similar fashion by manually selecting GO terms that represent pathway groups in *Drosophila.* Over-representation of categories was calculated using the associated genes of the identified regions as observation and all known genes in the genome or a specified region as background. The significance of the GO terms was calculated using R (http://www.r-project.org/) function ‘fisher.test’. *Drosophila* genome annotation (Version 5.1) was acquired from FlyBase (www.flybase.org).

### Interactive data mining tool

The enrichment analysis method presented in this paper is implemented as a user-friendly Shiny (Beeley, Chris. Web application development with R using Shiny. Packt Publishing Ltd, 2016) web-application accessible via http://gizmo2.cos.uni-heidelberg.de:3838. The user can select the set of genes to perform the GO enrichment analysis and the respective background independently. Results of the analysis are presented as a plot, an interactive table displaying significantly enriched GO groups, and an interactive heatmap, showing the counts of enriched GO terms within the respective higher-order GO group. It is also possible to get an insight into the individual GO terms that make up a category and into the genes that contributed to the categories or terms. The functionality of the tool exceeds what is described here, a detailed documentation of the tool is deposited under http://gizmo2.cos.uni-heidelberg.de:3838/documentation.html and additionally an interactive guide is provided in the online applicaiotn. After the login (username: damid17 and password: run) the user can access to the lists of genes active in soma (Soma) and genes bound by Abd-A or Zfh-1 or can upload his/her own data and analyze them using the tool.

### Gene interaction networks

The interaction networks were created using Cytoscape (http://www.cytoscape.org), an online tool that creates gene networks based on predicted, genetic, physical and shared protein domains interactions.

## AUTHOR CONTRIBUTIONS

F.P. conceived, designed, performed and interpreted experiments on Abd-A and the nucleoporins, worked together and supervised J.M., N.R., S.M., established the DamID protocols and wrote parts of the paper; S.T. conceived, designed, performed and interpreted experiments on V-ATPases, TOR and the transcriptional regulators, performed DamID experiments for Zfh-1, and soma and germline PolII TaDa, worked with E.R and O.E. on the interpretation of the bioinformatics data and wrote parts of the paper; E.R, O.E. and N.T. designed and implemented the integrative data mining and analysis tool, assembled with the help of F.P. and S.T. the list of higher-order GO terms; J.M. performed the DamID for Abd-A under the supervision of F.P.; J.Z. conceived, designed, performed and interpreted experiments on RNAi based analysis in the *Drosophila* intestine, M.B. conceived RNAi based analysis in the *Drosophila* intestine, J.U.L. assisted in designing and implementing the bioinformatics tool, I.L. conceived the study, assisted in designing and interpreting experiments, designed with E.R, O.E, N.T. and J.U.L. the data mining and analysis tool, wrote the paper, obtained funding to support the study (DFG/SFB 873).

## ACKNOWLEDGMENTS

We would like to thank the *Drosophila* community for providing us generously with fly stocks and antibodies, in particular Dorothea Goft for the TJ antibody, Ruth Lehmann for the Zfh-1 antibody, Margaret Fuller for the *c587-GAL4* fly lines, the Developmental Studies Hybridoma Bank (Iowa State University), the Vienna *Drosophila* Resource Center (VDRC), the Bloomington *Drosophila* Stock Center, the NIKON Imaging Center at Heidelberg University and the Genomics Core Facility at EMBL, Heidelberg. We would like to deeply thank M. Fuller, as part of the characterization of the Abd-A phenotype was done in her lab. We also thank all the members of the Ingrid Lohmann lab for critically reading the manuscript. We apologize to all whose work was not cited due to space limitations. This work was supported by the DFG/SFB 873, and an EMBO Short-Term Fellowship (ASTF 363 – 2012) to F. Papagiannouli to visit Stanford University.

